# BindingSiteAugmentedDTA: Enabling A Next-Generation Pipeline for Interpretable Prediction Models in Drug-Repurposing

**DOI:** 10.1101/2022.08.30.505897

**Authors:** Niloofar Yousefi, Mehdi Yazdani-Jahromi, Aida Tayebi, Elayaraja Kolanthai, Craig J. Neal, Tanumoy Banerjee, Agnivo Gosai, Ganesh Balasubramanian, Sudipta Seal, Ozlem Ozmen Garibay

## Abstract

While research into Drug-Target Interaction (DTI) prediction is fairly mature, generalizability and interpretability are not always addressed in the existing works in this field. In this paper, we propose a deep learning-based framework, called BindingSite-AugmentedDTA, which improves Drug-Target Affinity (DTA) predictions by reducing the search space of potential binding sites of the protein, thus making the binding affinity prediction more efficient and accurate. Our BindingSite-AugmentedDTA is highly generalizable as it can be integrated with any DL-based regression model, while it significantly improves their prediction performance. Also, unlike many existing models, our model is highly interpretable due to its architecture and self-attention mechanism, which can provide a deeper understanding of its underlying prediction mechanism by mapping attention weights back to protein binding sites. The computational results confirm that our framework can enhance the prediction performance of seven state-of-the-art DTA prediction algorithms in terms of 4 widely used evaluation metrics, including Concordance Index (CI), Mean Squared Error (MSE), modified squared correlation coefficient 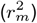, and the Area Under the Precision Curve (AUPC). We also contribute to the two most commonly used DTA benchmark datasets, namely Kiba and Davis, by including additional information on 3D structure of all proteins contained in these two datasets. We manually extracted this information from Protein Data Bank (PDB) files of proteins available at https://www.uniprot.org/. Furthermore, we experimentally validate the practical potential of our proposed framework through in-lab experiments. We measure the binding interaction between several drug candidate compounds for the inhibition of binding between (SARS-CoV-2 S-protein RBD) Spike and ACE-2 (host cell binding target) proteins. We then compare the computationally-predicted results against the ones experimentally-observed in the laboratory. The relatively high agreement between computationally-predicted and experimentally-observed binding interactions supports the potential of our framework as the next-generation pipeline for prediction models in drug repurposing.

## 1 Introduction

One critical step in drug designing and development is the identification of novel Drug-Target Interactions (DTIs), which characterize the binding of innovative candidate drug compounds to particular protein targets. Such experiments are often highly automated with chemical libraries for test compounds screened for activity in high-throughput methodologies. However, considering the enormous chemical and proteomic spaces, identifying novel compounds with specific interactive characteristics toward a particular protein target(s), if feasible, is often expensive, time- and resource-consuming Cohen (2002); Noble *et al*. (2004). To relieve this bottleneck, the use of computational methods is urgent to narrow down the search space for novel DTIs by predicting or estimating the interaction (strength) of novel drug–target pairs. A large body of studies proposed computational methods for DTI prediction tasks. Among these computational methods, docking simulations and machine learning methods are the two main approaches for *in-silico* prediction of DTI. Despite the accuracy and good interpretability of simulation and molecular docking, Salsbury Jr (2010) in virtual screening tasks, the performance of these structure-based computational methods highly depends on the high-resolution 3D structure of protein data and their availability. Additionally, these methods typically require tremendous computational resources, which tends to be a bottleneck for computational speed and prevents any large-scale applications of the methods. To overcome some of these hurdles, Artificial intelligence (AI), including Machine Learning (ML) and Deep Learning (DL) algorithms, have been introduced and applied across different stages of drug development and design pipelines. These models generally work by translating available knowledge about known drugs and their targets into some features, then used to train models that can predict interactions between new drugs or new targets. Earlier traditional machine learning methods use shallow models such as support vector machines, logistic regression, random forest, and shallow neural networks Yu *et al*. (2012); Tabei and Yamanishi (2013); Shi *et al*. (2013); Cheng *et al*. (2016) to address the problem of DTI prediction/estimation. However, these shallow models are often inadequate in capturing crucial features needed in modeling of complex interactions. Recently and powered by the increase in data and high-capacity computing machines, Deep Learning (DL) models have gained increased attention to address diverse problems in bioinformatics/cheminformatics applications, including drug discovery, especially DTI prediction tasks. DL approaches advance traditional shallow ML models due to their ability to automatically learn and extract feature representation, therefore identifying, processing, and extrapolating complex hidden interactions between drugs and targets Lee *et al*. (2019); Öztürk *et al*. (2018); Karimi *et al*. (2019); Öztürk *et al*. (2019). Since the focus of this study is Deep Learning models, which have gained much momentum over the last decades, we dedicate the entire Section 3 to discussing the state-of-the-arts that use DL approaches to address the problem of DTI prediction. In this study, we propose a computational-experimental framework that enables the next-generation pipeline for developing generalizable, interpretable, and more accurate prediction models in drug repurposing applications. At the core of our framework is an augmented-DTA ***in-silico*** prediction module, which utilizes a Graph Convolutional Neural Network (GCNN)-based model called AttentionSiteDTI that acts as a detector to identify the most probable binding sites of the target protein. This critical information is then augmented into a DL-based DTA prediction model to predict the binding affinities between drug-target pairs to enhance their prediction performance by narrowing down the search space of binding sites to the most promising ones. This approach is significant due to depending on the protein binding sites that are particularly sensitive and can be used to identify the ligands’ proper binding interactions HajiEbrahimi *et al*. (2017). As the computational results show, our framework leads to significantly improved performance of many state-of-the-art DTA prediction models in multiple evaluation metrics. We further design an ***in-vitro*** validation module, where we validate the practical application of our framework by comparing the computationally-predicted DTA values with those experimentally observed (measured) in the laboratory for several candidate compounds interacting with several target proteins. Also, our in-lab experimental validation illustrates improved agreement between computationally-predicted and experimentally-observed binding affinities between candidate compounds and proteins. Encouraged by the computational and experimental results, we then utilize our framework to accelerate the process of hit compounds identification in ***drug repurposing*** with the case study of SARS-Cov-2. Visualization of the proposed computational-experimental framework is illustrated in Figure 1.

**Fig. 1:**
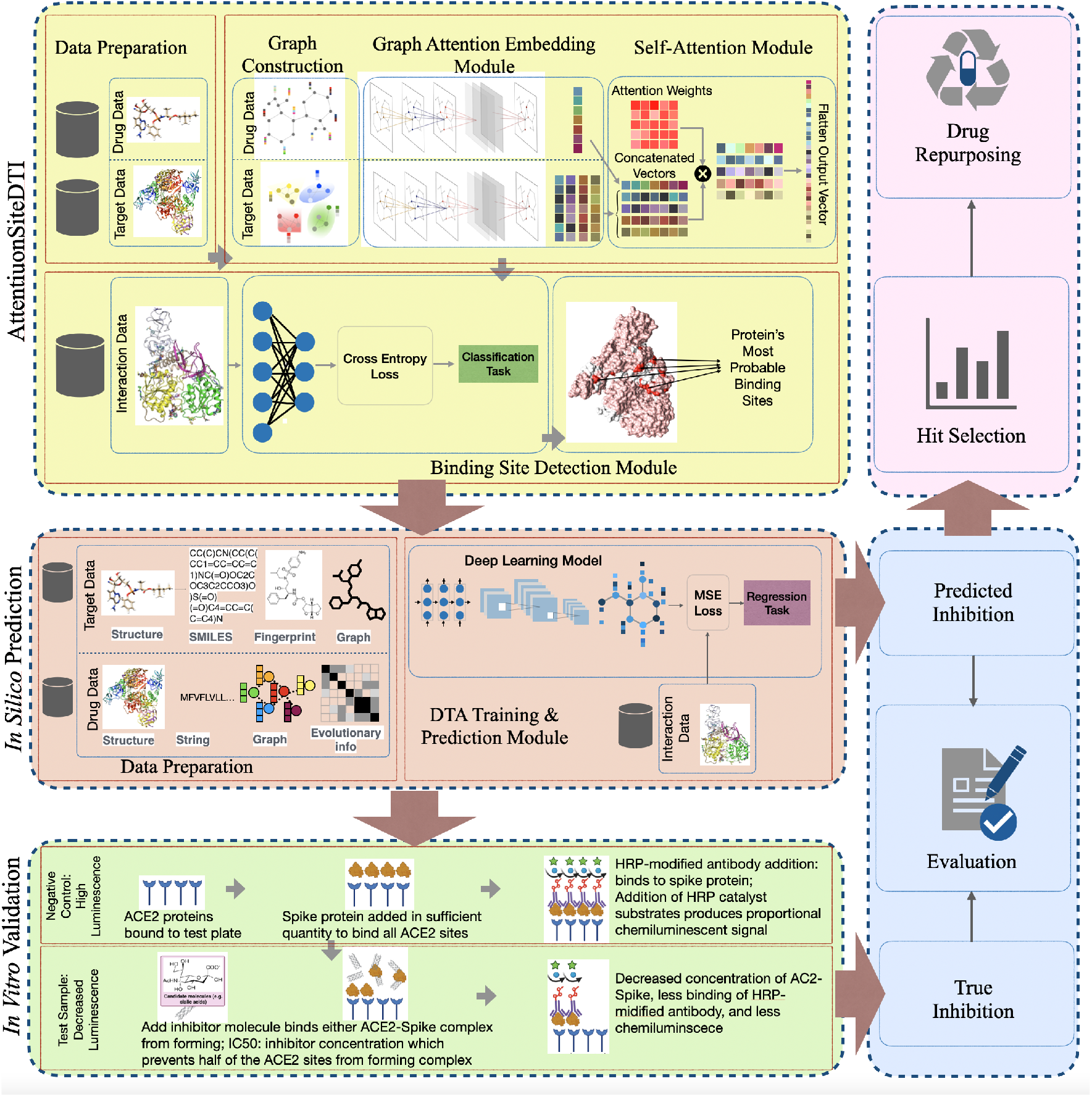
Our proposed framework includes three main modules: (1) *in-silico* prediction that consists of a DL-based drug-target affinity prediction model augmented with AttentionSiteDTI for finding the most probable binding sites of proteins; (2) *in-vitro* validation, where we compare our computationally-predicted results with experimentally-measured drug-target affinity values in laboratory to test and validate the practical potential of our proposed framework, and (3) drug repurposing module that utilizes our prediction model to identify hit compounds in prioritizing the most potent interactions for further *in-vitro* or *ex-vivo* verification in the laboratory.

### 1.1 Research Motivation

Over the last decade, most of the computational methods regarded DTI prediction as a binary classification problem, where the goal is to determine whether or not a drug–target (DT) pair interacts Ezzat *et al*. (2019). These methods ignore a vitally significant piece of information, namely the DT binding affinity values, reflecting the *strength* of interaction in drug-target pairs Thafar *et al*. (2019). Although several methods attend to the unbalanced nature of the DTI datasets Tayebi *et al*. (2022), predicting drug target affinity has many other implications; This motivates the use of regression approaches that are more informative and realistic but also more challenging in learning the strength of the binding between a drug and its target, which results in degraded prediction performance. Formulating the DTI prediction as a binding affinity regression task offers several advantages. First, it leads to more realistic prediction results, which reflect how tightly the drug ligand binds to a particular target protein; This, in turn, provides broader insight into many important aspects of the drug–target interactions, including their dose-dependence and drug efficacy, as well as discovering off-targets that can cause undesirable side effects. Second, predicting an approximate value for the strength of the interaction between the drug and target helps better with narrowing down the prospective drug candidates to be investigated via in-lab experiments that are expensive and time-consuming. The third advantage is the creation of a more realistic dataset, where DT pairs with no known binding information (missing or undiscovered) are not treated as negative (not-binding) samples. As argued in Wang *et al*. (2020b); Rifaioglu *et al*. (2020); Hu *et al*. (2019), the selection of non-binding interactions has a direct effect on model performance. Therefore, a regressing setting can not only better address the inherent complexity of the prediction task in the practical applications but also it can be easily transformed to either binary classification by setting specific thresholds or to ranking problem He *et al*. (2017), which offers more generalizability in practice. Although there exist approaches that treat the problem of DTI prediction using a regression framework, we argue that the performance of these regression models can be improved using a more efficient approach, wherein we first narrow down the search space by identifying the most probable binding sites of the protein in a drug-target pair, and then use this auxiliary piece of information in prediction of binding affinity values. To achieve this, we introduce a novel deep learning-based framework, BindingSite-AugmentedDTA, which not only improves the generalization and interpretation capabilities of the DTA prediction task but also leads to enhanced performance of many state-of-the-art models.

Despite the significant efforts that have been devoted to the development of computational models in recent years, their prediction power is rarely evaluated through in-lab experiments, which makes their practical benefits unknown, and thus limits their potential application in drug discovery and repurposing. Therefore, it is highly imperative to assess the prediction performance of *in-silico* models through experimental investigations in the laboratory in order to evaluate their practical power and to ensure their reliability in real-world predictions.

### 1.2 Contribution

The main contributions of our study are summarized as follows.

- We build an experimental-computational pipeline based on a graph convolutional neural network model, which is computationally and experimentally proven to lead to ***improved prediction performance*** when integrated with many state-of-the-art DL-based DTA prediction models. Also, our framework benefits from high ***generalizability*** and ***interpretability*** in prediction of drug-target affinity values:
  1. Our model is shown to significantly improve the prediction performance of many state-of-the-art models in terms of several performance metrics. The ***boost in the performance*** of these models is achieved by making them selectively focus on the most relevant parts of the input proteins when learning interaction relationships in drug-target pairs.
  2. Our model is highly ***generalizable*** because of to two main reasons: first is that it can be easily integrated with any DL-based prediction model, and the second is due to our target protein input representation that uses protein pockets (i.e., molecular fragments of binding sites influencing binding character) encoded as graphs. This helps the model concentrate on learning generic topological features from protein pockets, which can be generalized to new proteins that are not similar to the ones in the training data.
  3. Our model is highly ***interpretable*** in terms of the language of protein binding sites. Our model’s self-attention mechanism enables interpretability by understanding the prediction mechanism behind the model. In essence, this interpretability is enabled through learning which parts of the protein are more relevant in interacting with a given ligand. This is especially important in designing and developing new pharmaceutically active molecules, where it is crucial to know which parts of a molecule are essential for its biological properties.

- We demonstrate the prediction power of our model in practical applications through conducting ***in-vitro experiments***. To achieve this, we compare the computationally-predicted binding affinities (between some candidate compounds and a target protein) against the experimentally-measured binding affinities for the corresponding pairs. This is one of the main contributions of this work, as most of the *in silico* prediction models lack experimental validation, which is the ultimate determinant in assessing the validity of a computational model. Molecular dynamics (MD) simulations are additionally used as another computational yardstick to compare the in-lab experiments besides Deep Learning.
- Finally, inspired by the good performance of our proposed framework, we utilize it for ***drug repurposing*** by predicting binding affinity values between FDA-approved drugs and key proteins of SARS-CoV-2 including spike, 3C-like protease, RNA-dependent RNA polymerase(RdRp), helicase as well as Spike/ACE2 complex. We then provide a reference list of the top-ranked FDA-approved antiviral drugs with good binding affinities targeting different proteins of SARS-CoV-2.
- As a part of our work, we also contribute to the two most widely used benchmarking datasets in the DTA prediction task, namely Kiba and Davis datasets. These two datasets lack information on the 3D structure of proteins, which might be required by many advanced prediction models, including ours. We manually extract this information from Protein Data Bank (PDB) files of proteins available at https://www.uniprot.org/, and we provide the community with more complete versions of these two datasets.

## 2 BindingSite-AugmentedDTA for In-Silico Prediction

We utilize our recently developed model, called AttentionSiteDTI Yazdani-Jahromi *et al*. (2022), to improve the prediction performance of state-of-the-art DTA regression models. Our model finds the probability with which each binding site (pocket) of the protein will interact with the drug. Finding the most probable binding sites of the protein, we can then integrate this critical auxiliary information with DTA prediction models to enhance their performance. We refer to this customized version as BindingSite-AugmentedDTA, which can utilize any DL-based regression model to perform *Drug-Target Affinity* (DTA) prediction task. Given the fact that intermolecular interactions between protein and ligand occur in pocket-like regions of the protein (binding sites), identification of the most probable binding sites can help narrow down the search space, which, in turn, leads to more efficient and accurate DTA predictions.

As the computational and experimental results confirm, our model not only improves the prediction performance of state-of-the-art models, but also our self-attention bidirectional Long Short-Term Memory (LSTM) mechanism is proven to be useful in capturing the desired interaction features, which provides interpretability to the predictions. Moreover, instead of the standard interaction classification, we focus on a regression problem, where the task is to predict binding affinity values, reflecting the strength of the interaction in drug-target pairs. The results illustrate that our regression-based framework offers a more realistic, interpretable, yet accurate formulation of the drug-target interaction prediction task in practical applications. A brief description of our model is provided in Section 1.2, and our detailed approach is documented in Yazdani-Jahromi *et al*. (2022).

### 2.1 AttentionSiteDTI

Our proposed framework is built based on our previously developed model, called AttentionSiteDTI, which was initially developed for the classification task of Drug-Target Interaction (DIT) prediction. AttentionSiteDTI is inspired by models developed for sentence classification in the field of Natural Language Processing (NLP), where the drug-target complex is treated as a natural language sentence with structural and relational meaning between its biochemical entities, a.k.a. protein pockets and drug molecule. In this regard, each protein pocket or drug is analogous to a word, and each drug-target pair is analogous to a sentence. AttentionSiteDTI utilized an end-to-end Graph Convolutional Neural Network (GCNN)-based model to simultaneously learn context-sensitive graph embeddings of protein pockets and small molecules as well as a DTI prediction model capturing contextual and relational information contained in the sentence.

Unlike most of the graph-based models that use amino acid sequence representations for proteins, our model uses the 3D representation of pocket-like regions of the proteins as the input for target proteins. Considering the fact that the intermolecular interactions between protein and many ligands occur at different binding pockets (of the protein’s surface) rather than the whole protein, in our model, we represent protein pockets as graphs where the key protein residues correspond to the nodes that are connected based on residue proximity. Furthermore, the features associated with each node are encoded as a vector describing the local amino acid environment.

AttentionSiteDTI is highly generalizable due to the use of protein pockets encoded as graphs to represent the target protein. This allows the model to focus on learning generic topological features from protein pockets, which can be generalized to new proteins that are not similar to the ones in the training data. AttntionSiteDTI is also highly explainable due to its self-attention mechanism, which is used to capture any relationship between binding sites of a given protein and the drug in a sequence (i.e., sentence) and thus provide a better understanding of their binding relationships. To be self-contained, here we provide a brief description of AttentionSiteDTI, and we refer the readers to the original paper for more details Yazdani-Jahromi *et al*. (2022).

AttentionSiteDTI is an end-to-end graph-based deep learning model which was originally designed to address the problem of drug-target interaction prediction. It consists of four modules: (1) data preparation to find the binding sites of the proteins using the algorithm proposed in Saberi Fathi and Tuszynski (2014), (2) graph embedding learning module to learn the embeddings from constructed graphs of protein pockets and ligands as the inputs to the graph convolutional neural network, (3) prediction module to predict drug-target interactions using learned drug-target complex representations, and (4) interpretation module, quipped with a self-attention mechanism, to detect the most probable binding sites of the protein when interacting with the ligand in a given drug-target pair. The output of this module is indeed the main component that we use in this study to make the DTA prediction models focus on the most important parts of the protein when learning interactions between drug-target pairs. Figure 2 provides visualizations of our attention-based interpretation module where it detects the most probable binding sites of the main proteins of SARS-CoV2 in interaction with the drug named Remdesivir. Also, the overall architecture of our AttentionSiteDTI is shown in Figure 3.

**Fig. 2:**
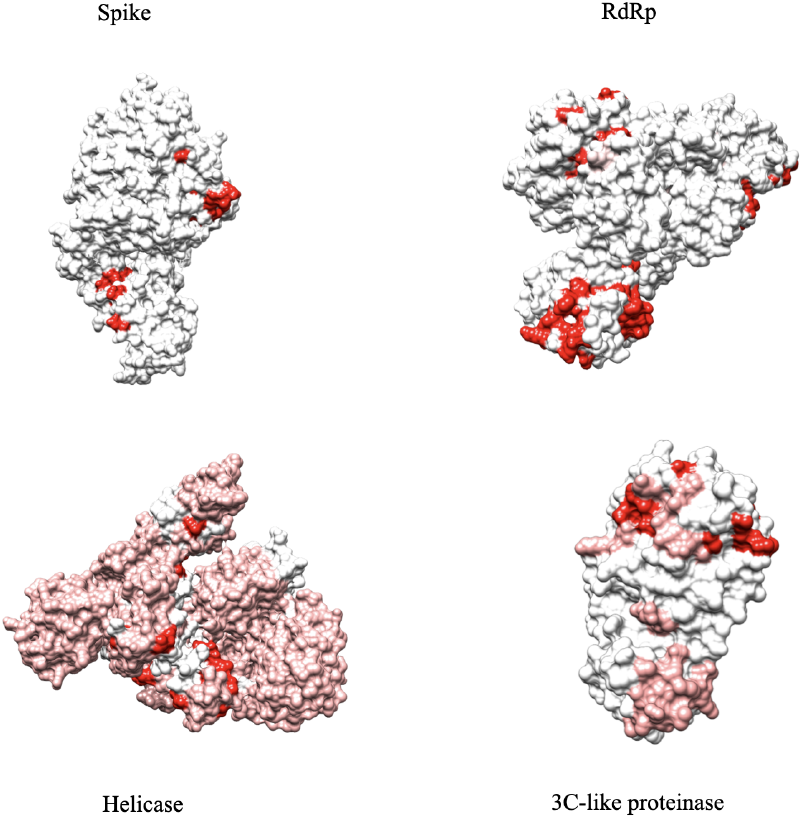
Pojected heatmap of self-attention weights on the 4 main proteins of SARS-Cov2. This figure shows the interpretability of our model, which can give us the binding site that has the most probability of binding to the ligand.

**Fig. 3:**
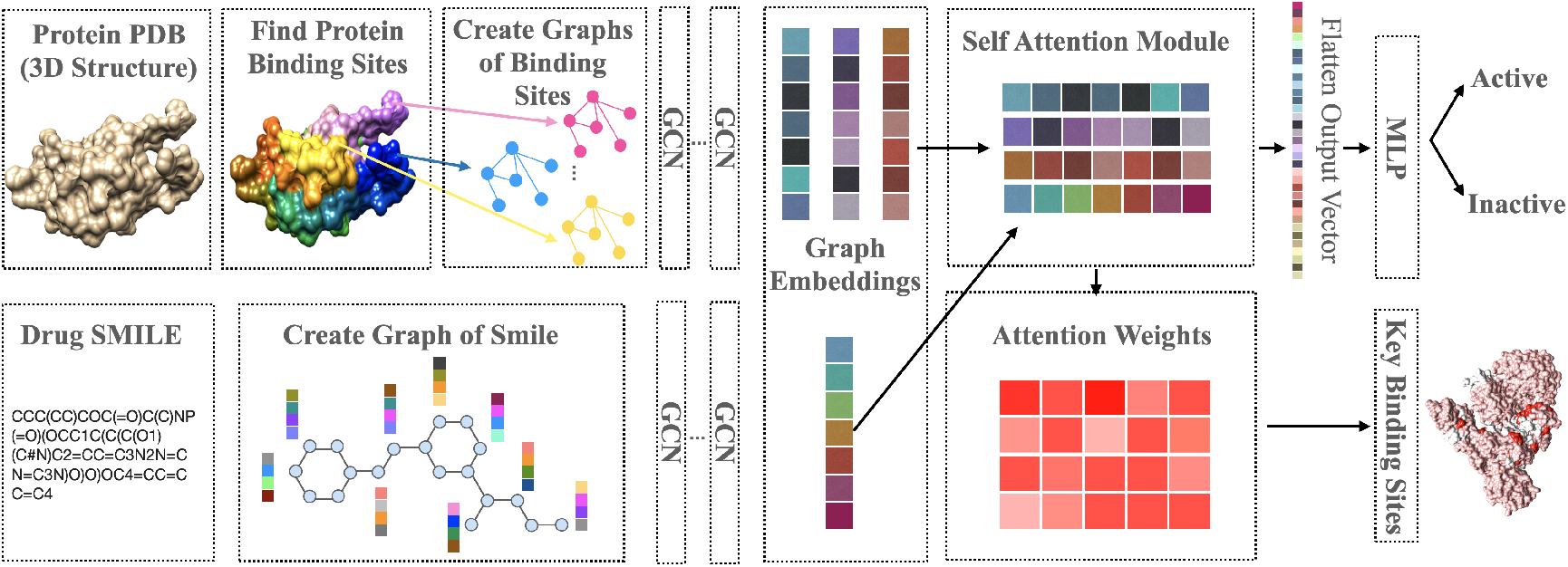
Architecture of AttentionSiteDTI: Following the extraction of protein’s binding sites, the graphs of protein pockets and ligands are constructed and fed into a graph convolutional neural network to learn corresponding graph embeddings. The concatenated representations are then fed into a binary classifier for predicting drug-target interactions. The self-attention mechanism in the network uses concatenated embeddings to compute the attention output, which enables interpretability by making the model learn the most probable binding sites of the protein in interaction with the ligand in a given drug-target pair.

## 3 Related Work: Deep Learning For Drug-Target Affinity Prediction

Inspired by the significant success of deep neural networks in the computer version, speech recognition, and NLP, they are now being widely used to address diverse problems in bioinformatics/cheminformatics applications Ekins (2016); Kalkatawi *et al*. (2019); Li *et al*. (2019), and specifically drug discovery Chen *et al*. (2018); Jing *et al*. (2018); Ekins *et al*. (2019). DL methods have been gaining popularity because of their efficiency and effectiveness in capturing low dimensional feature representations from simple sequence data, such as protein sequence and drug SMILES (Simplified Molecular Input Line Entry System), in less time and without requiring domain expertise. Indeed, these powerful methods can not only work in an end-to-end manner without any need to feature engineering but also they enable learning the hidden patterns in the data, and thus shown to outperform traditional ML models such as random forest (RF) Li *et al*. (2015); Shi *et al*. (2019); Zhao *et al*. (2021), support vector machines (SVM) Yu *et al*. (2012); Tabei *et al*. (2012); Mahmud *et al*. (2020) and other similarity rule-based approaches Fokoue *et al*. (2016).

Motivated by the success of the first studies that employed DL methods to model DTI, there has been an increasing number of later studies that adopted new DL architectures, such as convolutional neural networks (CNNs) and recurrent neural networks (RNNs) Gómez-Bombarelli *et al*. (2018); Jastrzę bski *et al*. (2016) as well as stacked-autoencoders Wang *et al*. (2018) to perform DTI binary interaction prediction using different input models for proteins and drugs.

Apart from different types of neural networks, various representations of the input data for the drug and target have been applied to train DL models for DTI prediction. These models can be categorized into two main branches in terms of data representation. First is sequence representation-based approaches, which take as input the sequence information of drugs and targets (e.g., SMILES for drugs and amino acid sequence for proteins). The second category includes those approaches with graph representations for drug and/or protein (e.g., graph representations for drugs and amino acid sequence for proteins).

A major limitation of the former is that these models use string representations for the drug compounds. However, this is not a natural and effective way to characterize molecules. Among all these approaches, we can refer to **DeepDTA** as the first approach that was introduced by Öztürk *et al*. (2018) to predict binding affinity as opposed to a binary (interact, no-interact) value. This model uses SMILES to represent drug input data and amino acid sequences to represent protein input data. Also, Integer/label encoding was used to encode both drug SMILES and the protein sequences. A CNN model with three 1D convolutional layers followed by a max-pooling function (called the first CNN block) was then applied to each drug embedding. The latest features for each protein are also learned using an identical CNN block. The feature vectors for each drug-target pair are then concatenated and fed into three FC layers and finally regressed with the drug–target affinity scores. **WideDTA** is an attempt to improve upon DeepDTA, which was introduced by the same authors Öztürk *et al*. (2019). Along with SMILES and amino acid sequences, WideDTA uses two other text-based information sources, which are Ligand Maximum Common Substructure (LMCS) for drugs, and Protein Domains and Motifs (PDM) based on PROSITE for proteins. Unlike DeepDTA, this model is a word-based model. That being said, both ligand SMILES and protein (amino acid) sequences are represented using a set of words instead of characters. More specifically, they represent a word in a drug SMILES using eight residues in the sequence and a protein with three residues in the sequence. A CNN model with four identical feature extracting blocks (extracting features from each of the text-based information sources) is then used to predict binding affinity scores. The concatenated features are then fed into 3 FC layers with two dropout layers to avoid overfitting. The output of the network is the scores for binding affinity for DT pairs. **AttentionDTA** Zhao *et al*. (2019) is another deep learning model that has been developed to predict compound–protein affinity using the semantic information of the drug’s SMILES string and protein’s amino acid sequence. This model uses two separate one-dimensional Convolution Neural Networks along with four different attention mechanisms to explore the relationship between drug and protein features.

As mentioned earlier, these sequence representations-based models fail at capturing the structural information of molecules, which in turn leads to the degraded predictive performance of the predictive model. On the other hand, a more natural way to describe the molecules seems to be graph representation, wherein the atom can be viewed as the nodes of the graph and chemical bonds as the edges. This form of data representation has inspired the use of DL models such as Graph Convolutional Neural Networks (GCNNs) that take graph structures as well as the characteristics of nodes or edges as their input data. Although GCNNs have already been used in drug discovery applications, including DTI, they are mainly focused on binary prediction task Gao *et al*. (2018); Mayr *et al*. (2018,?) to address the problem of DTI. That being said, among the graph-based approaches that frame the prediction problem as a regression task, we can refer to **GraphDTA** Nguyen *et al*. (2021) that predicts a continuous value for binding affinity. GraphDTA utilizes a GCCN-based module (with four different variants) on the molecular graph, describing drug molecules, along with a CCN-based module with three 1D convolutional layers, followed by a max pooling layer to extract feature representation of the input protein sequence. Finally, the concatenated vector of two representations is fed into several fully connected layers to predict the output DTA values. Inspired by GraphDTA, **DGraphDTA** Jiang *et al*. (2020) was also proposed for DTA prediction task. Similar to GraphDTA, DGraphDTA also uses graph representations for drug molecules. However, unlike GraphDTA, they use graph representation for protein sequences, as well. The focus of DGraphDTA (double graph DTA predictor) is on the protein graph representations. The protein graph construction is based on two sources of information. First is the protein residues as the nodes of the graphs; the Second is a representation called a contact map, which provides the spatial information of the interaction of residue pairs in the protein. More specifically, a contact map is a 2D (two-dimensional) representation of the 3D (three-dimensional) protein structure, usually in the form of a matrix consistent with the adjacency matrix in GNNs. The authors in Abdel-Basset *et al*. (2020) proposed a deep heterogeneous learning framework called **DeepH-DTA**, wherein three different modules have been used to extract features for drug molecules and protein sequences. Specifically, the drug molecules are encoded using two modules; one uses the SMILES representations along with a bidirectional ConvLSTM to model spatio-sequential information of the molecules, and the other one takes the topological drug information as input to generate drug representation using a Heterogeneous Graph Attention (HGAT) network. The third module takes the amino acid sequence of a protein to learn an embedding using a squeezed-excited dense convolutional network. The output of these three modules is finally concatenated to estimate the final prediction score of DTA. **DeepGS** Lin (2020) in another graph-based approach, which consider both local chemical context and the molecular structure in DTA prediction task. With the help of some embedding techniques such as Smi2Vec and Prot2Vec, DeepGS encodes the amino acid sequences of proteins as well as the atoms of drug molecules to distributed representations. Similar to deep-DTA, DeepGS learns two representations for the drug using two different modules; one uses the molecular graph as the input to a graph attention network (GAT) that extracts the topological information of the drug, and the second one takes an atom matrix A of embedding vectors (based on a pre-trained dictionary) as the input to a 1-layer bi-directional gated recurrent unit (BiGRU) to captures the local chemical context of atoms in the drug. A similar process to the second module of drug encoding was used to encode proteins using a CNN that allows capturing local chemical information of the amino acids. Finally, the concatenated vector of the three latent representations is fed into a stack of fully connected layers to predict the binding affinity scores for DT pairs.

## 4 Experiments and results

### 4.1 Experiments Setup and Hyperparameters

The hyper-parameter settings for the eight models were found by grid search and utilized and kept the same as reported in their original studies. For that, five training sets were used in 5-fold cross-validation to train the model, and the final CI scores were reported as the average of these five results. These hyper-parameters are summarized in Table 1 and Table 2.

**Table 1.**
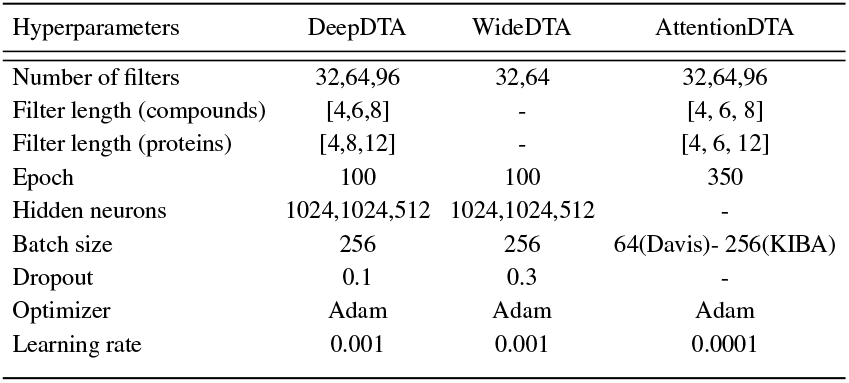
Summary of Parameter Setting for String Representation-Based Models

| Hyperparameters | DeepDTA | WideDTA | AttentionDTA |
| --- | --- | --- | --- |
| Number of filters | 32,64,96 | 32,64 | 32,64,96 |
| Filter length (compounds) | [4,6,8] | - | [4, 6, 8] |
| Filter length (proteins) | [4,8,12] | - | [4, 6, 12] |
| Epoch | 100 | 100 | 350 |
| Hidden neurons | 1024,1024,512 | 1024,1024,512 | - |
| Batch size | 256 | 256 | 64(Davis)- 256(KIBA) |
| Dropout | 0.1 | 0.3 | - |
| Optimizer | Adam | Adam | Adam |
| Learning rate | 0.001 | 0.001 | 0.0001 |

**Table 2.**
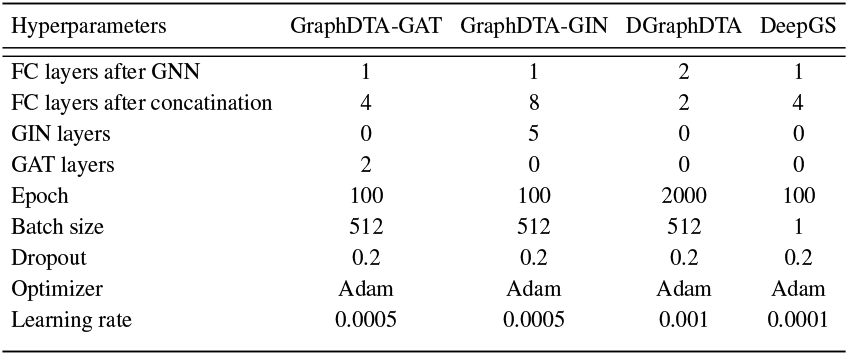
Summary of Parameter Setting for Graph Representation-Based Models

### 4.2 Benchmark Datasets

In this paper, two broadly used benchmark datasets for DTA, namely Davis Davis *et al*. (2011), and KIBA Tang *et al*. (2014) datasets, are used to compare the performance of our proposed model with the state-of-the-art models.

The Davis dataset contains 442 proteins related to the kinase protein family along their inhibitors (68 compounds) and the dissociation constant (*k*_*d*_) values corresponding to each assay. Similar to Öztürk *et al*. (2018); He *et al*. (2017) we convert the *k*_*d*_ values to log space, *pK*_*d*_ as explained in Equation 1

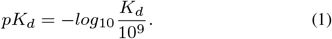

KIBA dataset combines different kinase inhibitor bioactivities (*k*_*i*_, *k*_*d*_ and IC50) to construct the KIBA score to optimize the consistency. He *et al*. (2017) filtered this dataset to include the drugs and targets with at least ten interactions, generating 229 unique proteins and 2111 unique drugs. Table 3 provides the statistics of the two datasets.

**Table 3.** Benchmark Datasets

| Dataset | Compounds | Proteins | Interactions |
| --- | --- | --- | --- |
| **KIBA** | 442 | 68 | 30056 |
| **Davis** | 229 | 2111 | 118254 |

### 4.3 Data Collection and Preprocessing

For the input of our model, we needed to extract PDB structures for the proteins. First, we extracted the identifier numbers for the proteins from https://www.uniprot.org/. Specifically, we restricted our search to human proteins, from which we chose the ones identified via the X-ray method. Although, there were a few proteins (put the number here, and maybe provide the complete info in the supplementary material.) for which no experimental data (PDB structure) were available. For these proteins, we used the PDB structures predicted by Alphafold (https://alphafold.ebi.ac.uk/).

### 4.4 Evaluation Metrics

Four widely used metrics in regression tasks were used to evaluate the performance of the models. Concordance index (CI) was first introduced by Gönen and Heller (2005) and is a ranking metric that measures whether the predicted binding affinity value of two random drug-target pairs is in the same order as their corresponding true values or not. CI metric is defined as:

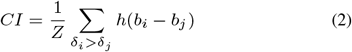

where *b*_*i*_ is the prediction value for the larger affinity *δ*_*i*_, and *b*_*j*_ is the prediction value for the smaller affinity *δ*_*j*_, *Z* is a normalization constant, *h*(*x*) is the step function Davies (2002); Pahikkala *et al*. (2015) which is defined as:

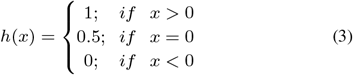

Mean Squared Error (MSE) Wackerly *et al*. (2014) is a commonly used loss function in regression-based models that measures the average squared difference between the predicted values and the actual values, and it is defined as:

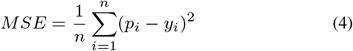

where *p*_*i*_ corresponds to the prediction value, and *y*_*i*_ represents the actual output. Also, *n* is the number of samples.

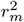 index Roy and Roy (2008) is another reported metric that was previously used to evaluate the external predictive performance of the models. The higher values (higher than 0.5) of this index determine whether a model is acceptable or not. As described in Pratim Roy *et al*. (2009); Roy *et al*. (2013), the metric is defined as below where *r*^2^ and 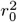 are the squared correlation coefficients with and without intercept:

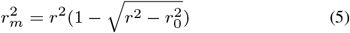

The Area Under Precision-Recall (AUPR) is another reported metric previously used in many studies. In order to measure this metric, the quantitative values were converted into binary values. Similar to Tang *et al*. (2014); He *et al*. (2017); Öztürk *et al*. (2018), the thresholds of *pK*_*d*_=7 for Davis dataset and 12.1 for KIBA dataset were used to utilize binarization.

### 4.5 Results and Comparison

To assess the effectiveness of our proposed framework, we compare the predictive power of six cutting-edge binding affinity DL-based prediction approaches with and without their integration with our AttentionSiteDTI module. The benchmarking approaches include both sequence representation-based models such as DeepDTA, WideDTA, and AttentionDTA, as well as graph representation-based approaches such as GraphDTA, DGraphDTA, and DeepGS. We compare the performance of all these models with and without the help of our AttentionSiteDTI in finding the most probable binding sites of the proteins. All models’ performances are evaluated under the same experimental conditions on the two benchmark datasets, KIBA and Davis. We used Concordance Index (CI), Mean Squared Error (MSE), modified squared correlation coefficient 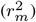, and the Area Under the Precision Curve (AUPC) for model evaluation. As the results in Figure 4 and Figure 5 illustrate, all six models show improved performance in our proposed regression framework when assisted with the AttentionSiteDTI model. The boost in models’ performance can be explained by the effectiveness of our model in finding the most probable binding sites of the proteins, which helps narrow down the search space of the prediction models by making them selectively concentrate on valuable parts of the proteins when learning the interactions between proteins and drugs; hence, enhancing the quality of the predictions. The missing values on the performance of the DeepH-DTA Abdel-Basset *et al*. (2020) is due to the missing information in the implementation of this model that was not provided in their code. Therefore, we were not able to produce the results of this model when combined with our AttentionSiteDTI. We gathered the results from the original paper on DeepH-DTA, as it was shown to outperform all other approaches in terms of all four metrics. However, its performance becomes very competitive with AttentionSite-augmented DGraphDTA and AttentionDTA on Kiba and Davis datasets, respectively.

**Fig. 4:**
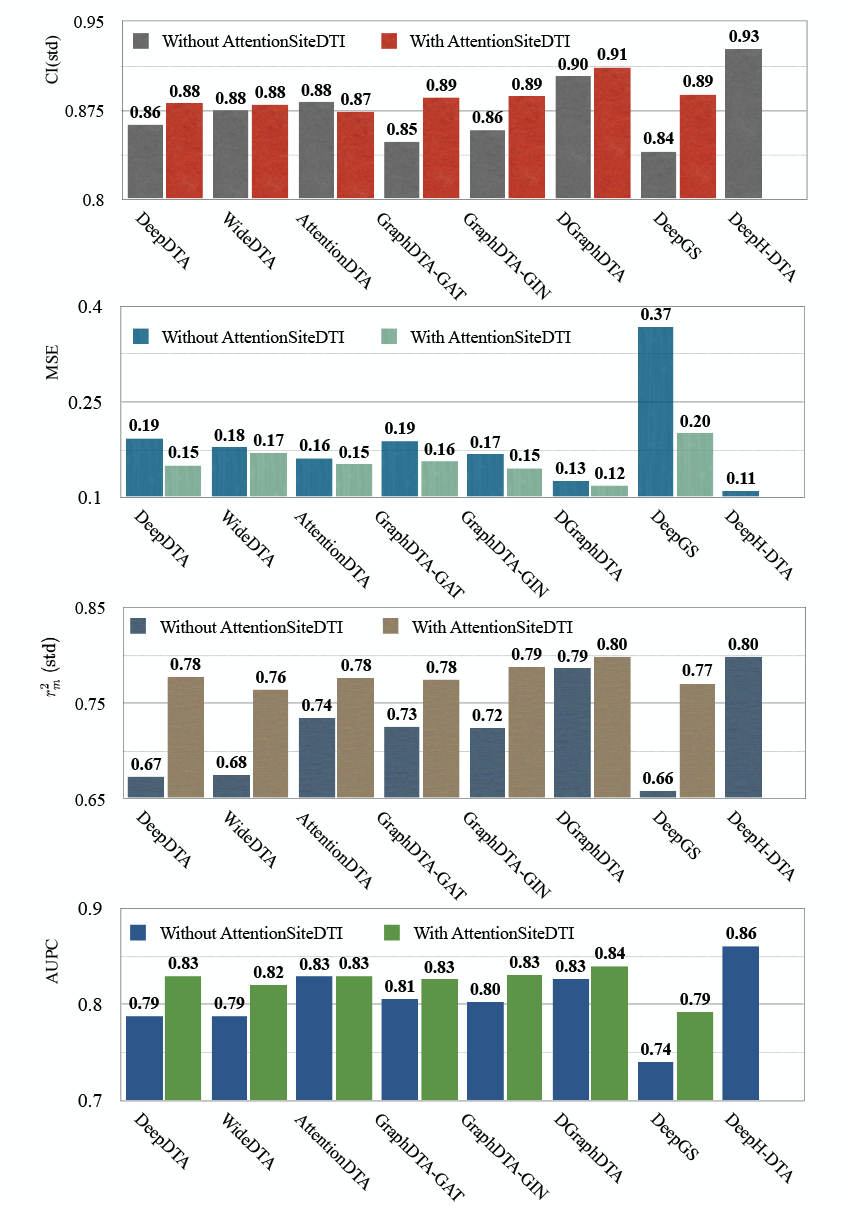
Performance of various DL-based DTA prediction models on KIBA dataset, with and without their integration with AttentionSiteDTI. Note that the scores of integrated DeepH-DTA do not show in the bar plots because of the missing information in the implementation of this model.

**Fig. 5:**
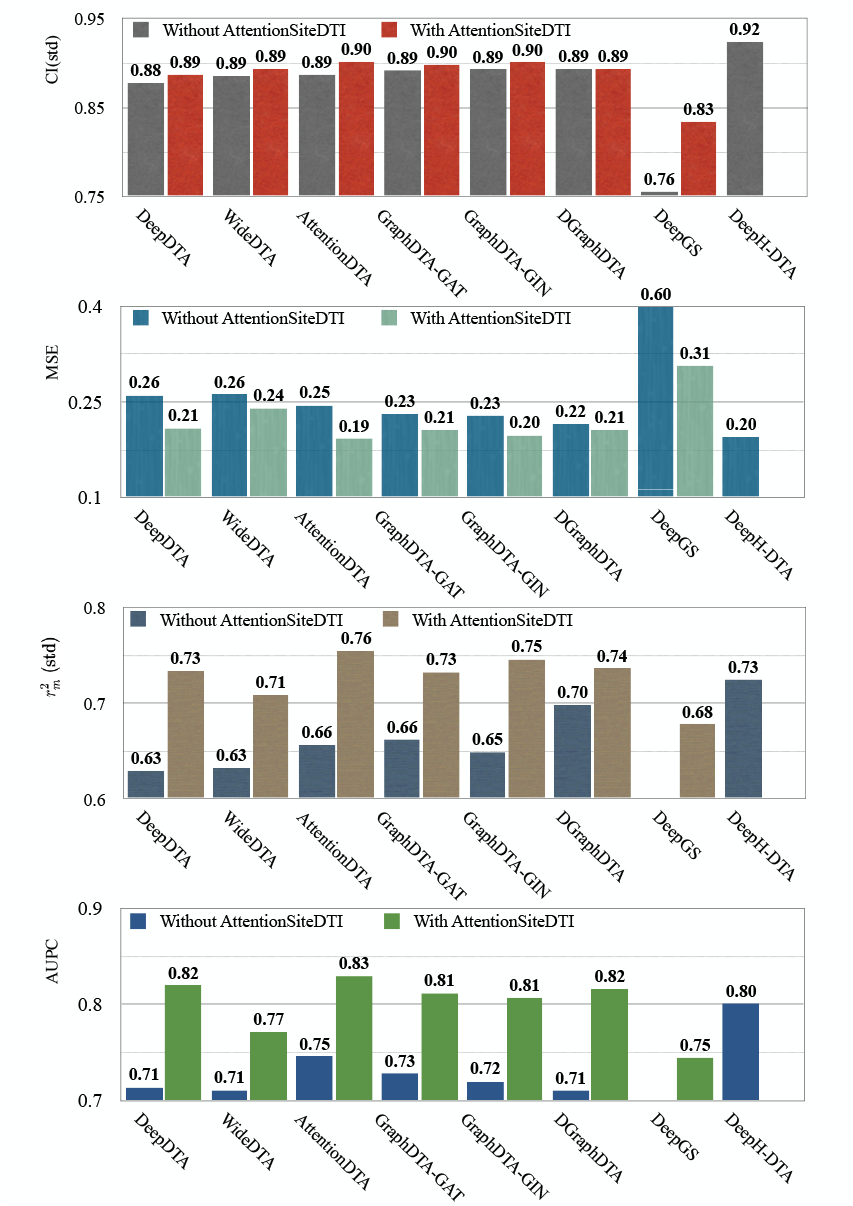
Performance of various DL-based DTA prediction models on Davis dataset, with and without their integration with AttentionSiteDTI. Note that the scores of integrated DeepH-DTA do not show in the bar plots because of the missing information in the implementation of this model. Also, the missing scores of DeepGS in the last two plots are because they are lower than the lower bound of the y-axis.

According to the results in Table 4, in the KIBA dataset, all AttentionSite-augmented models exceed their own plain versions in all evaluation metrics, where DeepGS enjoys the most improvement in all metrics, achieving 0.048, 0.166, 0.112 and 0.052 improvements for CI, MSE, 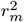, and AUPC, respectively. An exception is AttentionDTA, whose performance in CI metric in the augmented version is lower 0.008 than vanilla AttentionDTA. Among all metrics, 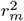 and *AUPC* have the most noticeable changes, where their normalized average improvements are 0.55 and 0.51 over all models. Note that we used min-max normalized of the difference scores to avoid the dependence on the choice of measurement units.

**Table 4.** Difference scores on performance metrics between vanilla and augmented models for KIBA dataset. The values in the table reflect *augmented*_*score*_ *− vanilla*_*score*_ for different performance measures. To calculate average values, we used min-max normalized of the difference scores to avoid the dependence on the choice of measurement units.

| Methods | Difference Scores |  |  |  |
| --- | --- | --- | --- | --- |
| | CI(std) | MSE | $r_m^2$ (std) | AUPC |
| String Representation-Based Approaches |  |  |  |  |
| DeepDTA | 0.018 | 0.043 | 0.105 | 0.042 |
| WideDTA | 0.005 | 0.009 | 0.089 | 0.033 |
| AttentionDTA | -0.008 | 0.01 | 0.042 | 0.001 |
| Graph Representation-Based Approaches |  |  |  |  |
| GraphDTA-GAT | 0.037 | 0.032 | 0.049 | 0.02 |
| GraphDTA-GIN | 0.028 | 0.021 | 0.064 | 0.028 |
| DGraphDTA | 0.008 | 0.007 | 0.013 | 0.013 |
| DeepGS | **0.048** | **0.166** | **0.112** | **0.052** |
| Min | -0.008 | 0.007 | 0.013 | 0.001 |
| Max | 0.048 | 0.166 | 0.112 | 0.052 |
| Normalized Average | 0.022 | 0.215 | 0.553 | 0.510 |

For the Davis dataset, as the results show in Table 5, the augmented models also achieve better results compared to their plain versions. similarly, DeepGS benefits the most from AttentionSiteDTI, where it attains 0.088 0.291, 0.438, 0.198 improvements for CI, MSE, 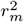, and AUPC, respectively. Again, the two performance metrics 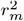 and *AUPC* show the most average changes with 0.23 and 0.32 improvements across all models.

**Table 5.** Difference scores on performance metrics between vanilla and augmented models for Davis dataset. The values in the table reflect *augmented*_*score*_ *− vanilla*_*score*_ for different performance measures. To calculate average values, we used min-max normalized of the difference scores to avoid the dependence on the choice of measurement units.

| Methods | Difference Scores |  |  |  |
| --- | --- | --- | --- | --- |
| | CI(std) | MSE | $r_m^2$ (std) | AUPC |
| String Representation-Based Approaches |  |  |  |  |
| DeepDTA | 0.009 | 0.052 | 0.104 | 0.106 |
| WideDTA | 0.008 | 0.021 | 0.076 | 0.061 |
| AttentionDTA | 0.015 | 0.053 | 0.098 | 0.084 |
| Graph Representation-Based Approaches |  |  |  |  |
| GraphDTA-GAT | 0.006 | 0.026 | 0.071 | 0.084 |
| GraphDTA-GIN | 0.008 | 0.031 | 0.097 | 0.087 |
| DGraphDTA | 0 | 0.009 | 0.038 | 0.116 |
| DeepGS | **0.088** | **0.291** | **0.438** | **0.198** |
| Min | 0 | 0.009 | 0.038 | 0.061 |
| Max | 0.088 | 0.291 | 0.438 | 0.198 |
| Normalized Average | 0.218 | 0.213 | 0.234 | 0.322 |

It is worth mentioning that there exist other DL-based models such as Cui *et al*. (2019); Xia *et al*. (2020); Stepniewska-Dziubinska *et al*. (2020); Mylonas *et al*. (2021); Kandel *et al*. (2021), which were specifically proposed for binding site prediction task. However, none of them have been shown to be generalizable in integration with DL-based DTA prediction models. Indeed, to the best of our knowledge, we are the first to propose this unified framework, which also provides interpretability to any integrated models by providing a deeper understanding of which binding sites of a target protein are most probable to bind with a given ligand. From the figures in these two tables, we can also observe that sequence-based DL models have lower predictive power compared to graph-based models, which explains the effectiveness of graph representations in capturing structural information of drugs and/or proteins as deciding factors in the prediction of DTIs.

## 5 In-Vitro Validation: SARS-Cov-2 Case study

To evaluate the practical potential of our proposed framework, we further experimentally tested and validated its performance through experimentations in the laboratory by measuring the binding affinities between 13 candidate compounds and the Spike-ACE2 complex. Additional information on our choice of compounds is provided in supplementary material. We then assessed the prediction performance using three metrics: (1) Pearson correlation coefficient to measure the scoring power, indicating the binding affinity prediction capacity of the model, (2) Concordance Index (CI) to measure the ranking power, presenting the affinity-ranking capacity of the model and (3) Mean Square Error (MSE) to measure the overall prediction error, showing the quality of the model in estimating the true values. To predict compound-protein binding affinities, we selected DeepDTA because of its wide adoption in previous studies as the first DL approach developed to predict drug-target binding affinity. DeepDTA is among state-of-the-art methods that have shown relatively good performance compared to many DL-based models with higher complexity in architecture and computation time. In order to show the effectiveness of our prediction framework, the performance measures were calculated under two settings; with and without the incorporation of AttentionSiteDTI with the DeepDTA regression model. Since the model should always be trained under distinct prediction scenarios and regarding the fact that the few labeled data points from lab experiments are inadequate for training the models from scratch, we train AttentionSitDTI and DeepDTA on the BindingDB dataset, which includes a wide variety of drug-like compounds and target proteins. This enables both models to learn generic rules governing the ligand-protein interactions from the BindingDB dataset, containing 2,303,972 binding data for 8,561 protein targets and 995,797 small molecules, as of July 2021. Once the *general* complex interaction patterns are extracted, the models then utilize this pre-learned knowledge and transfer it to the *specific* task of binding affinity prediction between 13 drug-like compounds and the Spike-ACE2 complex. These compounds include Darunavir, N-acetyl-neuraminic acid, N-Acetyllactosamine, 3*α*, 6*α* Mannopentaose, N-glycolylneuraminic acid, 2-Keto-3-deoxyoctonate ammonium salt, cytidine5-monophospho-N-acetylneuraminic acid sodium salt, Congo red, Direct Violet, Evans Blue, Calcomine scarlet 3B, Chlorazol Black and Methylene Blue as inhibitor molecules to bind to the Spike protein, or the ACE2 receptor protein (as the primary host factor recognized and targeted by SARS-CoV-2 Spike protein).

ML has been recently used to enhance the sampling of MD simulation trajectories to capture millisecond scale in-lab processes using nanosecond scale simulations. It has also been used to remove noise in simulation data and make it more accessible for interpretation Wang *et al*. (2020a); Ribeiro and Tiwary (2019). Free energies calculated from MD simulations have been used to train DL models to accelerate the prediction of free energies based on the structural information of small molecules Bennett *et al*. (2020). Joshi et al. Joshi *et al*. (2020) have used DL based prediction framework to screen molecules from the Selleck database and perform molecular docking to further optimize the search results for final use in MD simulations to find the molecules having the most potential to bind with the main protease of SARS-CoV-2. In the present study, a comparison has been made between MD simulations of certain selected molecules with the RBD of SARS-CoV-2 from the pool of molecules checked using DL-based frameworks and in-lab experiments. The purpose is to check the similarity of DL predictions with that of MD simulations and use a two-prong approach to guide in-lab experiments. The details are presented as supplementary information.

As the results in Table 6 indicate, the AttentionSiteDTI helps enhance the performance of DeepDTA, especially in the two metrics, MSE and Pearson correlation coefficient, where the improvements are 1.08 and 0.38, respectively. As a matter of fact, when using DeepDTA, we observe a very low correlation of 0.04 between the predicted and measured bioactivities, which is significantly boosted to 0.42, considered a relatively good correlation as a result of higher quality binding affinity predictions by *augmented* DeepDTA. In fact, this illustrates the effectiveness of our proposed framework in improving the performance of DeepDTA when augmented with our AttentionSiteDTI. Note that Figure 6 reports the predicted and measured bioactivity profiles of Spike-ACE2 complex against 13 drug-like compounds tested in our experimental assay.

**Table 6.** Performance comparison of the computational models on in-vitro experimental bioactivity results

| Methods | CI(std) | MSE | Pearson Correlation |
| --- | --- | --- | --- |
| DeepDTA | 0.4358 | 3.7729 | 0.0463 |
| DeepDTA + AttentionSiteDTI | **0.4487** | **2.6907** | **0.4245** |

**Fig. 6:**
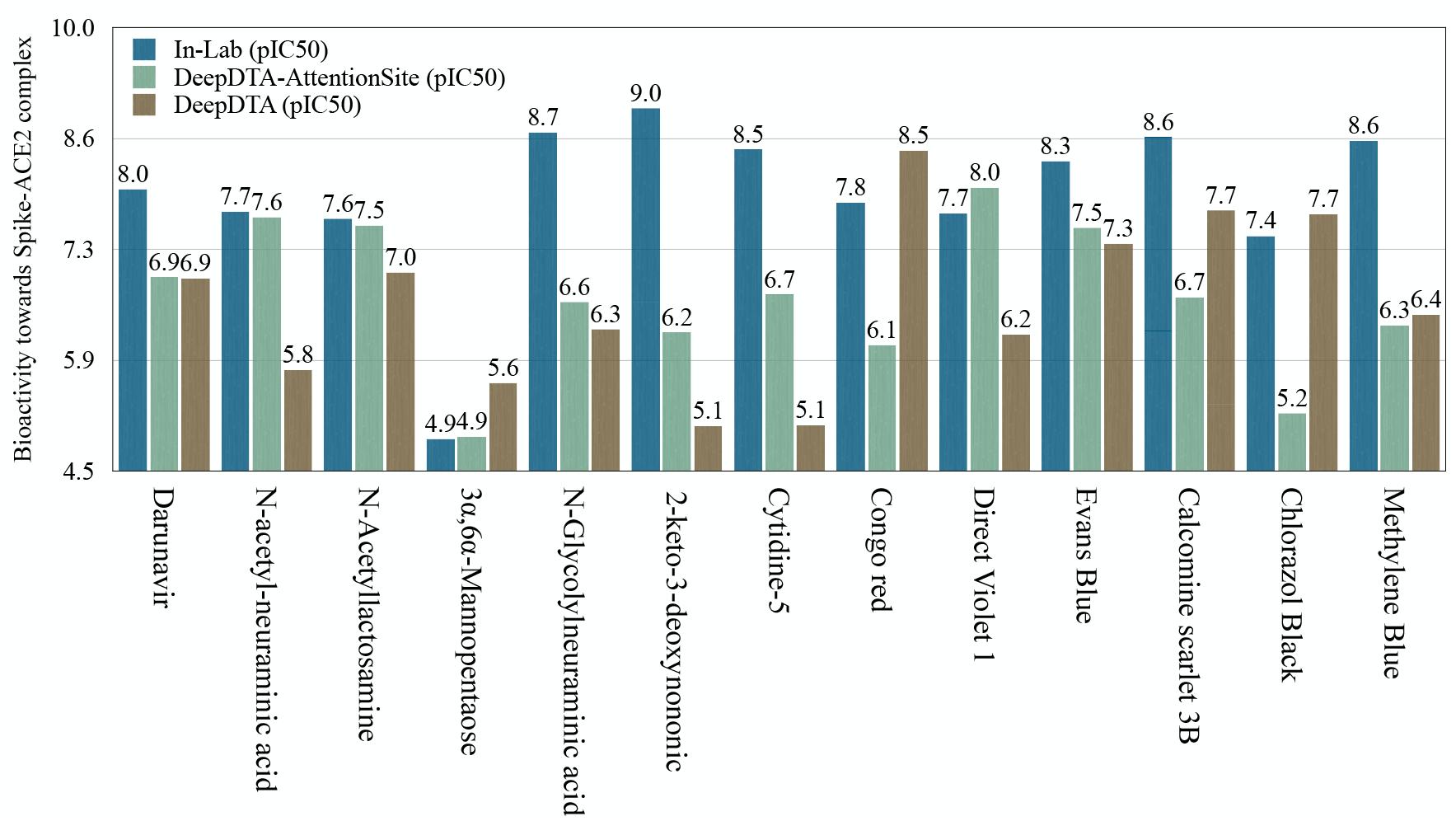
Predicted and measured binding affinity values of Spike-ACE2 complex with 13 drug-like compounds in our experimental assay. The predicted values are reported for DeepDTA model with and without its integration with AttentionSiteDTI.

For the third approach mentioned here, we have initially performed molecular docking using AutoDock Vina and AutoDock tools, and the highest scored ligand-protein complex conformation structures are chosen for the further all-atom Steered Molecular Dynamics (SMD) simulations. Here, only 5 of the previously mentioned 13 molecules have been simulated using SMD simulations to determine the peak unbinding force and center of mass separation of the corresponding ligands with time (Supplementary material Figures 3 and 4). Our molecular docking results show that three ligands bind to one single binding pocket of the protein, which is in line with previously mentioned theories of having specified ligand binding locations in spike protein (Supplementary material Section 5.1). Next, our SMD simulation results showed that N-Acetyl neuraminic acid (sialic acid) requires the highest amount of steering force before complete dissociation takes place (Supplementary material Figure 3 and Table 2). Based on this, the five molecules analyzed via SMD simulations are ranked in such a way that rank one is given to the molecule that shows the highest binding strength and force requirement, while the lowest ranked molecule has the least tendency to bind with the protein in a favorable environment. The comparison of this ranking analyzed via different in silico approaches, MD simulations, and in-lab experiments based on binding affinity is given in Supplementary information Table 3. From Table 7, it is clearly seen that both our SMD simulation and DeepDTA-AttentionSite model predict N-acetyl neuraminic acid as the highest ranked molecule in terms of binding affinity with spike protein, and interestingly, DeepDTA, our MD simulation, and the in-lab experiments all predict N-glycolyl neuraminic acid as the second based inhibitor. Cytidine-5’ has been ranked 5th both by DeepDTA and our SMD simulation as the molecule least prone to bind, but the N-acetyllactosamine is ranked last in the in-lab experiment. Therefore, even though there are similarities in results based on the DL model and our SMD simulations, the in-lab experiment results seem to vary for the molecules having the least binding affinity. So, the in-lab experiments might be optimized based on the DL and MD results to accelerate the selection of potential inhibitors.

**Table 7.** Top 10 FDA-approved antiviral drugs predicted by AttentionSite-Augmented DeepDTA model to have highest affinity scores with 4 genome sequences related to SARS-CoV-2

| Drug Name | PIC50 | Rank in 3416<br>FDA-Approved Drugs | Rank in 58<br>Antiviral Drugs |
| --- | --- | --- | --- |
| **Spike-ACE2** |  |  |  |
| Oseltamivir phosphate | 8.29 | 42 | 1 |
| Etravirine | 8.22 | 52 | 2 |
| Fosamprenavir calcium | 8.20 | 57 | 3 |
| Tipranavir | 7.94 | 108 | 4 |
| elvitegravir | 7.88 | 141 | 5 |
| ganciclovir | 7.84 | 162 | 6 |
| Simeprevir | 7.71 | 221 | 7 |
| Rimantadine | 7.67 | 248 | 8 |
| saquinavir | 7.65 | 259 | 9 |
| danoprevir | 7.57 | 314 | 10 |
| **helicase** |  |  |  |
| Fosamprenavir calcium (FPV) | 7.51 | 21 | 1 |
| Simeprevir | 7.47 | 24 | 2 |
| Elvitegravir | 7.28 | 52 | 3 |
| Simeprevir sodium | 7.11 | 87 | 4 |
| elvitegravir | 7.09 | 97 | 5 |
| ritonavir | 7.01 | 120 | 6 |
| Indinavir sulfate | 6.98 | 132 | 7 |
| Amprenavir | 6.93 | 155 | 8 |
| nelfinavir | 6.88 | 182 | 9 |
| Tenofovir alafenamide fumarate | 6.86 | 190 | 10 |
| **3C-like** |  |  |  |
| Fosamprenavir calcium | 7.72 | 5 | 1 |
| elvitegravir | 7.25 | 29 | 2 |
| Tipranavir | 7.17 | 37 | 3 |
| Asunaprevir | 7.08 | 46 | 4 |
| Elvitegravir | 7.06 | 47 | 5 |
| Atazanavir sulfate | 6.92 | 68 | 6 |
| Daclatasvir | 6.88 | 77 | 7 |
| Saquinavir mesylate | 6.77 | 97 | 8 |
| Simeprevir | 6.71 | 121 | 9 |
| Simeprevir sodium | 6.68 | 129 | 10 |
| **RdRp** |  |  |  |
| elvitegravir | 7.60 | 8 | 1 |
| Fosamprenavir calcium | 7.24 | 24 | 2 |
| ritonavir | 6.96 | 53 | 3 |
| Asunaprevir | 6.90 | 66 | 4 |
| Simeprevir sodium | 6.87 | 71 | 5 |
| Daclatasvir | 6.85 | 74 | 6 |
| Simeprevir | 6.82 | 79 | 7 |
| nelfinavir | 6.81 | 82 | 8 |
| Amprenavir | 6.79 | 93 | 9 |
| Saquinavir mesylate | 6.78 | 96 | 10 |

## 6 Drug Repurposing: SARS-Cov-2 Case study

Drug repurposing is the process of identifying new applications for existing drugs to treat new emerging diseases. Repurposing can help accelerate the identification of drugs that have already proven to be safe and effective in humans and are tested for potential adverse effects and interactions. It has become a promising approach due to the opportunity for reduced costs and timelines required for the development of new drugs. The greatest value of this expedited process becomes even more prominent when facing outbreaks of a novel, challenging infections, including the COVID-19 pandemic, which demands fast and cost-effective tools and techniques to identify novel treatments. Although repurposing can be applied at any stage of drug development, its most significant benefit is with already (FDA) approved drugs, as it can lead to notable savings with the regulatory process needed for approving new drugs. For example, the benefits of drugs such as Remdesivir, Favipiravir, Ribavirin, Lopinavir, Ritonavir, Arbidol, Darunavir, Tocilizumab, Interferons, and Dexamethasone have been noticed in different clinical studies to save the life of COVID-19 patients Chakraborty *et al*. (2021).

In the era of big data, the application of AI approaches to address the problem of drug repurposing is not only formidable but also necessary. To this end, there are four different AI-enabled approaches that can be used to tackle this problem Gupta *et al*. (2021). The first approach is based on biomedical knowledge graphs, which is a systematic way to incorporate expert-derived sources of information into a graph with nodes representing entities (such as proteins and drugs) and edges capturing relationships between those entities. The second approach is molecular docking, which is a simulation-based approach to predicting the interaction between some selected protein targets and ligand drugs. The third approach is known as gene expression signatures, which involves the comparison of gene expression signatures of new candidate therapies to those of known effective treatments. The fourth approach, which is also the focus of this study, is based on the idea of *Drug-Target Affinity (DTA)* prediction that we discussed earlier.

Encouraged by the excellent performance of our framework, we now apply it to predict binding affinity values between 3445 commercially available FDA-approved drugs (including 85 antiviral drugs) and key proteins of SARS-CoV-2, including 3C-like protease, RNA-dependent RNA polymerase(RdRp), helicase as well as Spike/ACE2 complex. We use AttentionSite-augmented DeepDTA, which is pre-trained on the BindingDB dataset to predict binding affinities between genome sequences of SARS-CoV-2 —extracted from the National Center for Biotechnology Information (NCBI) database—with 85 FDA-approved antiviral drugs from DrugBank. The exhaustive list of drugs is provided in the supplementary material.

As the results show in Table 7, the Spike protein–ACE2 interface was predicted to have the highest binding affinity with Oseltamivir phosphate, Etravirine and Fosamprenavir calcium, all with PIC50 values greater than 8 nM, which are followed by other antiviral drugs with predicted binding affinities of PIC50 > 7 nM potency (we only included top 10 antiviral drugs in Table 7 with highest binding affinity). Among the prediction results, Fosamprenavir calcium, Simeprevir, Elvitegravir, Simeprevir sodium, elvitegravir and ritonavir have shown good inhibition towards SARS-Cov-2 helicase, with predicted binding affinity values above 7 nM. Furthermore, Fosamprenavir calcium, elvitegravir, Tipranavir, Asunaprevir and Elvitegravir were predicted to have a potential affinity (PIC50> 7 nM) to SARS-Cov2 3C-like Proteinase. Finally, elvitegravir and Fosamprenavir calcium were the only inhibitors that were predicted to have binding affinities greater than 7 nM in binding with RNA-dependent RNA polymerase. These verifying studies are available in Table 7. Despite the existence of any strong real-world evidence on the effectiveness of these drugs (except for the FDA-approved drug Remdesivir) against COVID-19, we found that some of these candidate drugs have already been suggested or introduced by other studies, including in-silico, preclinical, and clinical trials.

A more interesting observation in our prediction results can be seen in Table 8. There are eight antiviral drugs that are currently in clinical trials to be assessed for their efficacy against COVID-19. These drugs include Atazanavir, Daclatasvir, Danoprevir, Darunavir, Elvitegravir, Lopinavir, Oseltamivir and Ritonavir, which are also listed in the present prediction results as suitable potential inhibitors to all four subunits of SARS-Cov-2 (Spike-ACE2 interface, helicase, 3C-like protease, and RdRp), mostly with binding affinity values greater than 6 nM in PIC50. Also, Remdesivir, which is the only FDA-approved drug for the treatment of COVID-19, shows great predicted potency to all subunits of SARS-Cov-2 as follows: against Spike-ACE interface (PIC50 7.47 nM), RNA-dependent RNA polymerase (PIC50 6 nM), helicase (PIC50 6 nM) and 3C-like protease(PIC50 5.68 nM).

**Table 8.** Binding affinity values of selected antiviral drugs in clinical trials to treat COVID-19, binding with different SARS-CoV-2 related receptors. ^* *^ The information on the number of clinical trials and supporting references are gathered from https://covid19-help.org/ as of July 2022.

| Drug Name | # Clinical Trial | # Supporting References | Spike-ACE2 |  | helicase |  | 3C-like |  | RdRp |  |
| --- | --- | --- | --- | --- | --- | --- | --- | --- | --- | --- |
|  |  |  | PIC50 | rank | PIC50 | rank | PIC50 | rank | PIC50 | rank |
| Atazanavir | 3 | 6 | 6.89 | 36 | 6.36 | 32 | 6.18 | 23 | 6.10 | 21 |
| Daclatasvir | 8 | 5 | 7.23 | 24 | 6.57 | 21 | 6.88 | 7 | 6.85 | 6 |
| Danoprevir | 2 | 3 | 7.57 | 10 | 5.68 | 54 | 6.46 | 17 | 5.71 | 33 |
| Darunavir | 2 | 6 | 6.40 | 50 | 6.72 | 15 | 5.85 | 39 | 6.05 | 24 |
| Elvitegravir | 7 | 2 | 7.43 | 14 | 7.28 | 3 | 7.06 | 5 | 6.61 | 12 |
| Lopinavir | 39 | 26 | 7.44 | 12 | 5.55 | 60 | 5.26 | 60 | 5.18 | 53 |
| Oseltamivir | 9 | 5 | 6.75 | 41 | 5.24 | 70 | 5.27 | 59 | 5.12 | 57 |
| **Remdesivir** | **67** | **70** | **7.47** | **11** | **6.00** | **44** | **5.68** | **43** | **6.00** | **26** |
| ritonavir | 57 | 27 | 7.16 | 26 | 7.01 | 6 | 6.62 | 11 | 6.96 | 3 |

## 7 Conclusion

In this paper, we proposed a framework that can enhance Drug-Target Affinity (DTA) predictions by first finding the most probable binding sites of the protein, thus, making the binding affinity prediction more efficient and accurate. Our AttentionalSiteDTA is not only highly generalizable as it can be combined with any DL-based regression model but also provides interpretability to integrated models due to its attention mechanism, which enables learning which binding sites of a protein interact with a given ligand. The computational results confirm that our framework leads to improved prediction performance of seven state-of-the-art DTA prediction algorithms in terms of 4 widely used evaluation metrics, including Concordance Index (CI), Mean Squared Error (MSE), the modified, squared coefficient of correlation 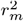, and the Area Under the Precision Curve (AUPC). Furthermore, we experimentally validated the prediction potential of our framework by conduction *in-vitro* experiments. To achieve this, we measured the binding affinities between 13 drug-like compounds and the Spike-ACE2 interface and compared these experimental results against binding affinity values that we predicted by our computational models. The results of our in-lab validation illustrated the potential of our framework to significantly enhance the performance of the widely-adopted DeepDTA model in the prediction of binding affinity values in real-world applications. Also, following the analysis of our drug screening, we proposed several FDA-approved antivirals that could display antiviral activities against SARS-Cov-2. Some of these compounds are currently undergoing clinical trials, and some others are also suggested by other studies for further evaluation in experimental assays and clinical trials to investigate their actual activity against COVID-19. Another contribution of this work is to provide the community with extended versions of the two most commonly used datasets, Kiba and Davis, for which we manually extracted information on the 3D structure of all proteins in these two datasets from the Protein Data Bank (PDB) files of proteins available in https://www.uniprot.org/.

## Data and Code availability

All datasets and codes are publicly available at https://github.com/yazdanimehdi/BindingSite-AugmentedDTA

## Competing interests

There is no competing interest.

## Notes

### Competing Interest Statement

The authors have declared no competing interest.

